# Release of acidic store calcium is required for effective priming of the NLRP3 inflammasome

**DOI:** 10.1101/2022.01.06.475262

**Authors:** Nick Platt, Dawn Shepherd, Yuzhe Weng, Grant C. Churchill, Antony Galione, Frances M. Platt

## Abstract

The lysosome is a dynamic signaling organelle that is critical for cell functioning. It is a regulated calcium store that can contribute to Ca^2+^-regulated processes via both local calcium release and more globally by influencing ER Ca^2+^ release. Here, we provide evidence from studies of an authentic mouse model of the lysosomal storage disease Niemann-Pick Type C (NPC) that has reduced lysosomal Ca^2+^ levels, and genetically modified mice in which the two-pore lysosomal Ca^2+^ release channel family are deleted that lysosomal Ca^2+^ signaling is required for normal pro-inflammatory responses. We demonstrate that production of the pro-inflammatory cytokine IL-1β via the NLRP3 inflammasome is significantly reduced in murine Niemann-Pick Type C, the inhibition is selective because secretion of TNFα is not diminished and it is a consequence of inefficient inflammasome priming. Synthesis of precursor ProIL-1β is significantly reduced in macrophages genetically deficient in the lysosomal protein Npc1, which is mutated in most clinical cases of NPC, and in wild type cells in which Npc1 activity is pharmacologically inhibited. Comparable reductions in ProIL-1β generation were measured *in vitro* and *in vivo* by macrophages (MΦ) genetically altered to lack expression of the two-pore lysosomal Ca^2+^ release channels *Tpcn1* or *Tpcn2*. These data demonstrate a requirement for lysosome-dependent Ca^2+^ signaling in the generation of specific pro-inflammatory responses.

## Introduction

Since their discovery by Christian de Duve (1955) lysosomes have been recognized as the major cellular site of macromolecular degradation and re-cycling^1^. However, more recent studies have revealed that the lysosome is a much more complex organelle that is involved in multiple additional cell biological processes^2–3^ including nutrient sensing^4^, energy metabolism^5^, cell repair^6^, cell death^7^, cell migration^8^, immunity^9^ and infection^10^. In particular, the lysosome has been found to be a dynamic signaling hub that integrates and delivers cellular signals from the environment and directly interacts and signals with other organelles. Fundamental to this was the discovery of the lysosome as a regulated cellular calcium (Ca^2+^) store, in addition to the endoplasmic reticulum^11,12^.

The importance of the lysosome in cellular health has been underscored by the study of a group of rare, inherited metabolic diseases, collectively known as lysosomal storage diseases (LSDs). At least seventy, predominantly autosomal recessive LSDs have been described, in which lysosomal dysfunction occurs^13^. Although there is a spectrum of clinical phenotypes both between and within specific diseases, a wide range of organs and tissues can be affected and most have a neurodegenerative clinical course^13^. A currently unexplained universal feature of LSDs is the induction of aberrant inflammation that actively contributes to disease pathogenesis^13,14^. Myeloid cell activation, particularly of microglia in the CNS, leads to elevated levels of multiple pro-inflammatory cytokines and chemokines, and the recruitment of inflammatory cell populations into affected tissues^15,16–19^.

Niemann-Pick disease Type C (NPC) is a progressive, neurodegenerative disorder in which multiple lipid species accumulate in the late endo-lysosomal system and many cell types are affected^20,13^. This disorder most frequently (95% of clinical cases) results from loss of function of the lysosomal membrane protein, NPC1^20^. Although the precise function of NPC1 remains unresolved, the disease is characterized by accumulation of sphingosine in the endo lysosome, which inhibits re-filling of the lysosomal with Ca^2+^, resulting in reduced lysosomal Ca^2+^ content and diminished Ca^2+^ release when mobilized by the second messenger, nicotinic acid adenine dinucleotide (NAADPs)^21,22^.

In this study, we report an unexpected pro-inflammatory profile in NPC that includes significantly impaired production of the prototypic cytokine, IL-1β. This cytokine is biologically potent and has extensive effects within inflammatory and immune responses^23–24^. IL-1β is produced by the coordinated action of cytosolic protein complexes known as inflammasomes^25^. Canonical generation of bioactive IL-1β is by the NLRP3 inflammasome, the most well-characterized inflammasome. This occurs by a two-step process involving an initial priming step, typically the result of binding of a pro-inflammatory ligand, such as lipopolysaccharide (LPS) or pro-inflammatory cytokine to cognate pattern recognition or cytokine receptor^25^. This priming step results in transcription and translation of ProIL-1β. A second activating signal is then required, which includes a wide range of stimuli such as adenosine triphosphate (ATP), pore-forming toxins and particulates that then induce inflammasome assembly, caspase-mediated cytokine maturation and secretion^26,27^.

Here, we show that priming of the NLRP3 inflammasome, which is responsible for the generation of ProIL-1β is perturbed in genetic and pharmacological models of NPC and that this phenotype is replicated in cells in which release of lysosomal Ca^2+^ via two pore channels (TPCNs) is pharmacologically or genetically blocked. These data reveal that lysosomal Ca^2+^ regulates the nature and extent of the pro-inflammatory cytokine response following challenge, specifically through its regulation of NLRP3 inflammasome priming.

## Results

### Resident peritoneal macrophages (RPMΦ) from Npc1^-/-^ mice have increased acidic compartment volume, size and granularity

Expansion of the late endosome/lysosome (LE/Lys) compartment in cells is a hallmark of LSDs ^28^. We made use of an authentic mouse model of NPC1, and performed FACS analysis of cells from a peritoneal lavage from symptomatic *Npc1*^-/-^ and age-matched *Npc1*^+/+^ mice stained with anti-F4/80 antibody and LysoTracker™-green DND-26. Gating on F4/80^hi^ cells (Suppl Fig 1), we confirmed that as predicted RPMΦ from *Npc1*^-/-^ mice had statistically higher LysoTracker™ staining, consistent with an increased relative acidic compartment volume (Fig 1). *Npc1*^-/-^ myeloid cells were also significantly larger (increased FSC-A) and displayed greater side scatter (SSC-A), a measure of intracellular complexity/granularity, than the equivalent population of myeloid cells from wild type mice. There was no significance difference in the fluorescence intensity of staining for the MΦ marker, F4/80 between *Npc1*^-/-^ and *Npc1*^+/+^ mice.

**Fig 1.**
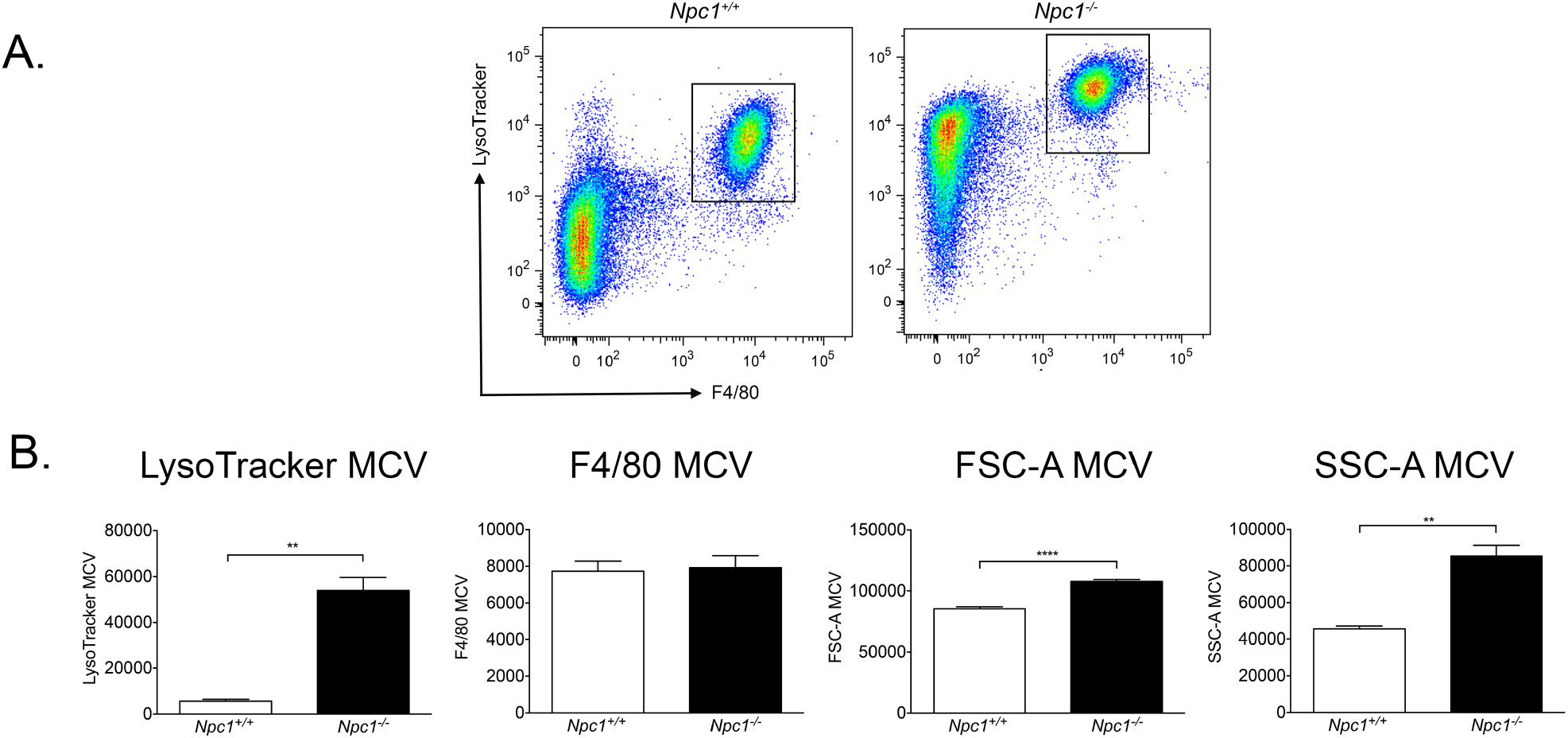
*Npc1*^-/-^ resident peritoneal macrophages (RPMΦ) display significantly greater LysoTracker™ staining, indicating expansion of relative acidic compartment volume. Panel A. Representative FACS profiles of *Npc1*^+/+^ (left) and *Npc1*^-/-^ (right) RPMΦ. Y-axis indicates relative LysoTracker™ staining intensity; x-axis indicates relative F4/80 staining intensity. Gated cells are F4/80^hi^ RPMΦ. Panel B. Histograms of relative intensity of LysoTracker™ fluorescence, relative intensity of F4/80 staining, relative FSC-A and relative SSC-A values for *Npc1*^+/+^ (open columns) and *Npc1*^-/-^ (filled columns) RPMΦ., ** *p*<0.01. Student *t*-test. Data are representative of three independent experiments.

### Npc1^-/-^ RPMΦ secrete significantly less IL-1β after stimulation but normal levels of TNFα

In order to investigate the production of IL-1β, we primed *Npc1*^+/+^ and *Npc1*^-/-^ RPMΦ with different concentrations of LPS, which is a ligand for Toll-like receptor 4, then activated the cells with ATP for 1h and collected culture supernatants, which were analyzed for cytokine content by specific ELISAs. We found significantly lower levels of IL-1β secreted by *Npc1*^-/-^ RPMΦ in comparison to *Npc1*^+/+^ RPMΦ at all concentrations of LA used for priming (Fig. 2A). We saw comparable reductions in IL-1β production when RPMΦ were primed with peptidoglycan before activation with ATP (Fig 2B), indicating that impairment of IL-1β production was independent of the nature of the priming stimulus. To determine whether the reduced secretion of IL-1β was a reflection of a generalized deficit in pro-inflammatory cytokine generation or restricted to this specific cytokine, we assayed TNFα in supernatants after appropriate stimulation. There was no significant difference between the amounts of TNFα produced by *Npc1*^+/+^ and *Npc1*^-/-^ RPMΦ (Fig 2C).

**Fig 2.**
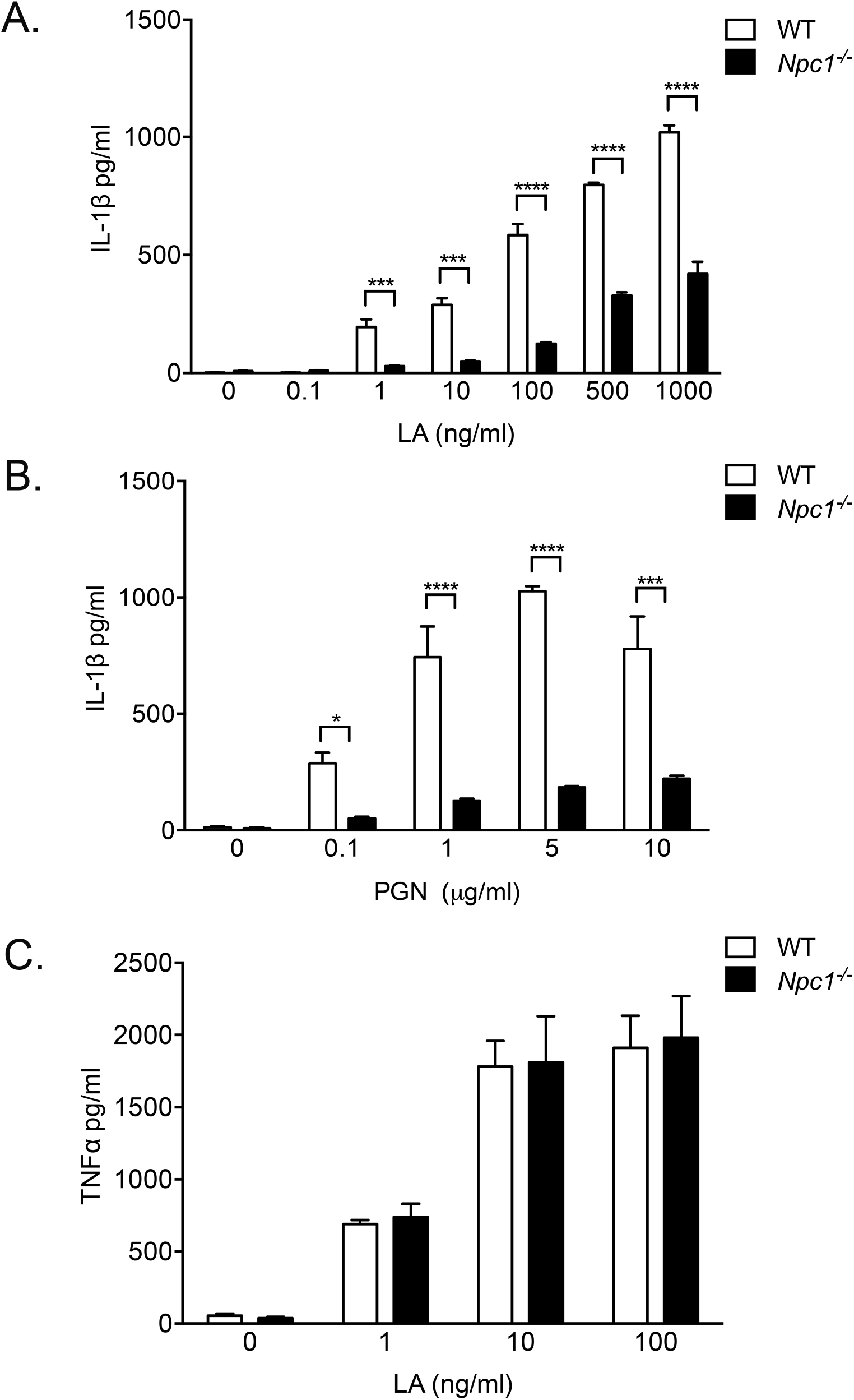
Primed and activated *Npc1*^-/-^ resident peritoneal macrophages (RPMΦ) secrete significantly less IL-1β than *Npc1*^+/+^, but equivalent amounts of TNFα. Panel A, IL-1β concentrations (pg/ml) in culture supernatants of *Npc1*^+/+^ (open columns) and *Npc1*^-/-^ (filled columns) RPMΦ primed with various concentrations of LA and then activated with 5 mM ATP. Panel B, IL-1β concentrations (pg/ml) in culture supernatants of *Npc1*^+/+^ (open columns) and *Npc1*^-/-^ (filled columns) RPMΦ primed with various concentrations of peptidoglycan and then activated with 5 mM ATP. Panel C, TNFα concentrations (pg/ml) in culture supernatants of *Npc1*^+/+^ (open columns) and *Npc1*^-/-^ (filled columns) RPMΦ stimulated with various concentrations of LA. Data shown are mean± SEM, n= 4 per treatment condition. **** *p*< 0.0001, *** *p*<0.001, ** *p*<0.01, * *p*<0.05. 2-way ANOVA. Data are representative of two independent experiments.

### Reduction in IL-1β generation by Npc1^-/-^ RPMΦ is a result of impaired priming of the NLRP3 inflammasome

To determine whether NLRP3 priming or activation or both steps are affected in NPC cells we used multiple methodological approaches to discriminate between these possibilities. We probed western blots of RPMΦ cell lysates and cell supernatants of cells that had been primed with lipid A (LA) or primed and activated with extracellular ATP. Lysates from LA-primed *Npc1*^-/-^ cells contained significantly less ProIL-1β p35 in comparison to wild type MΦ (Fig. 3A). As a consequence, activated *Npc1*^-/-^ deficient cells secreted significantly less mature cytokine (p17) than wild type RPMΦ.

**Fig 3.**
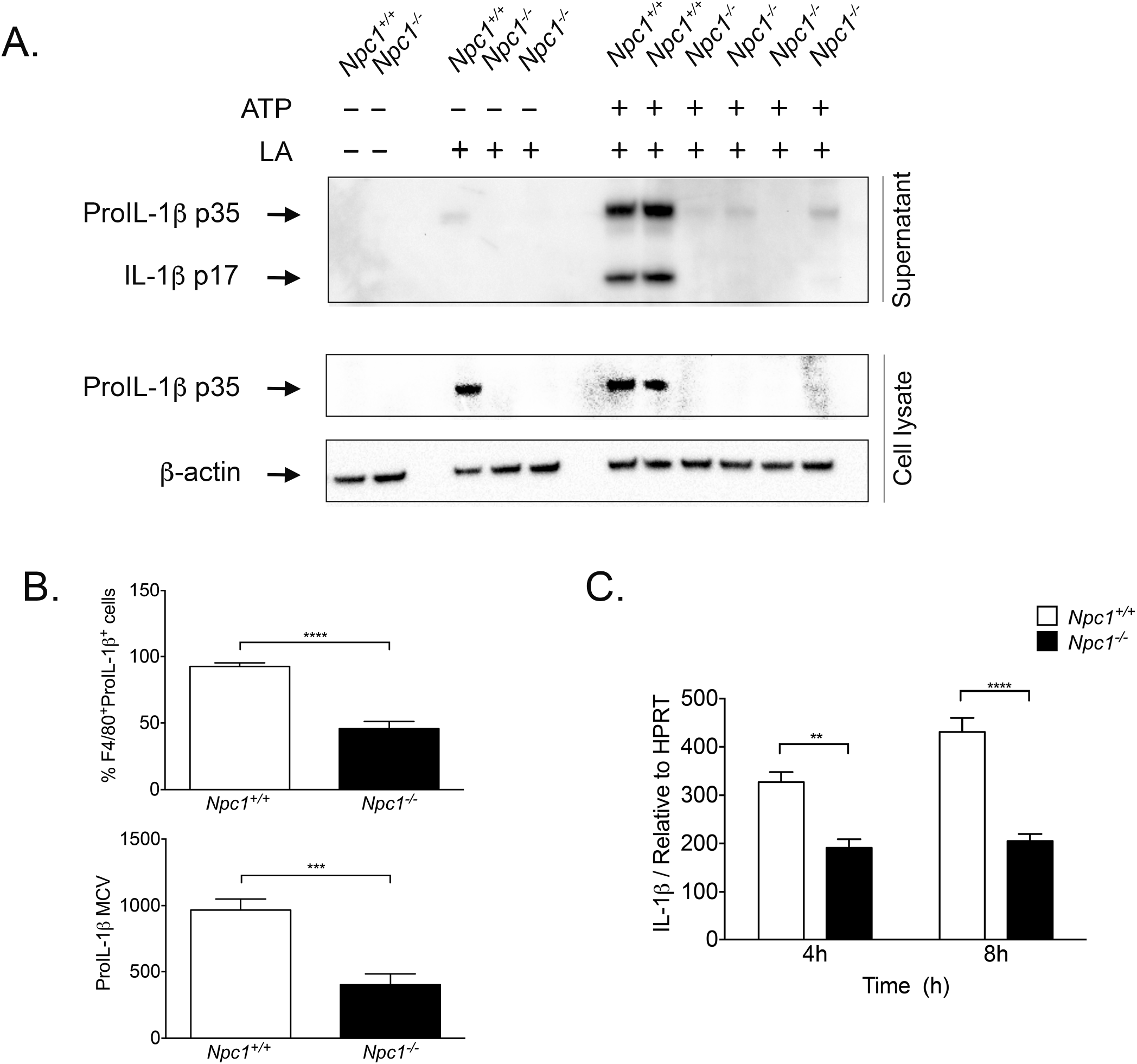
Significantly impaired priming of the NLRP3 inflammasome in *Npc1*^-/-^ RPMΦ. Panel A. Western blot of culture supernatant proteins and cell lysates of *Npc1*^+/+^ and *Npc1*^-/-^ RPMΦ untreated, primed with 100ng/ml LA and where indicated, activated with 5 mM ATP and probed with anti-IL-1β antibody. Bands corresponding to ProIL-1β p35 and IL-1β p17 proteins are indicated. Panel B. Histogram of the frequencies (upper) and relative staining intensities (lower) of *Npc1*^+/+^ (open columns) and *Npc1*^-/-^ (filled columns) RPMΦ primed with 100 ng/ml LA, stained with anti-ProIL-1β antibody and analyzed by FACS. MCV = mean channel value. Data are mean± SEM, n= 6. **** *p*< 0.0001, *** *p*<0.001. Student t-test. Panel C. Relative expression of IL-1β transcripts in *Npc1*^+/+^ (open columns) and *Npc1*^-/-^ (filled columns) RPMΦ after priming with 100 ng/ml LA for 4h or 6h. Data are mean± SEM, n= 6. **** *p*< 0.0001, ** *p*<0.01. 2-way ANOVA. Data are representative of two independent experiments.

In light of these data, we took advantage of an antibody (clone NJTEN3) that specifically recognizes murine ProIL-1β and does not bind to the mature cytokine to confirm reduced generation of the precursor cytokine by *Npc1*^-/-^ RPMΦ after priming. FACS analysis allowed us to determination not only the frequency of cytokine-positive cells but also the relative levels of ProIL-1β per cell. We observed a significantly reduced frequency of ProIL-1β^+^ F4/80^hi^ *Npc1*^-/-^ cells after LA stimulation, which had a significantly lower fluorescence intensity than the equivalent population of primed *Npc1*^+/+^ RPMΦ (Fig 3B).

To determine whether the lower levels of ProIL-1β in primed *Npc1*^-/-^ RPMΦ was the result of reduced transcription we used Q-PCR to determine relative transcript abundance. Data obtained were consistent with significantly lower ProIL-1β transcript abundance in stimulated *Npc1*^-/-^ RPMΦ (Fig 3C).

### Pharmacological inhibition of Npc1 significantly reduces generation of IL-1β by murine RAW 264.7 MΦ cells

U18666A is a drug that binds directly to Npc1^29^ and inhibits its function, inducing all of the cellular phenotypes that uniquely characterize this disease at the cellular level^21,30^. To exclude the possibility that impaired generation of IL-1β by primed *Npc1*^-/-^ RPMΦ might result from the influence of other peritoneal cell types and to enable temporal control of induction of NPC cellular phenotypes, we treated RAW 264.7 cells with U18666A before NLRP3 priming and activation. FACS analysis of LysoTracker™ stained cells confirmed U18666A drug treatment caused a significant increase in acidic compartment volume, consistent with induction of NPC cellular phenotypes (Suppl Fig 2). Western blotting of cell supernatants from U186666A treated cells that had been primed with increasing concentrations of LA and then activated with ATP revealed a significant decrease in the amounts of ProIL-1β p35 and mature IL-1β p17 released from U18666A-treated in comparison to vehicle-treated RAW 264.7 MΦ (Fig 4A). We made a similar investigation of the murine J774.1 MΦ cell line and obtained comparable results; a dose-dependent reduction in cytokine generation by U18666A-treated cells as compared to control (Suppl Fig 3). Quantification of bioactive IL-1β in RAW 264.7 MΦ culture supernatants by ELISA confirmed significantly lower cytokine secretion by drug-treated cells (Fig 4B). In contrast, there was a statistically significant increase in secretion of TNFα by RAW 264.7 MΦ stimulated with different doses of LA (Fig 4B). Q-PCR analysis of IL-1β transcription over a 24h time course following LA stimulation verified statistically lower transcription in U18666A-treated RAW 264.7 MΦ at 6, 8 and 24h time points (Fig 4C). There was a comparable reduction in *caspase-1* transcription at multiple time points analyzed, but there were no differences in transcription of *Nlrp3* and *Asc* genes between MΦ that had been either vehicle or U18666A treated (Fig 4C).

**Fig 4.**
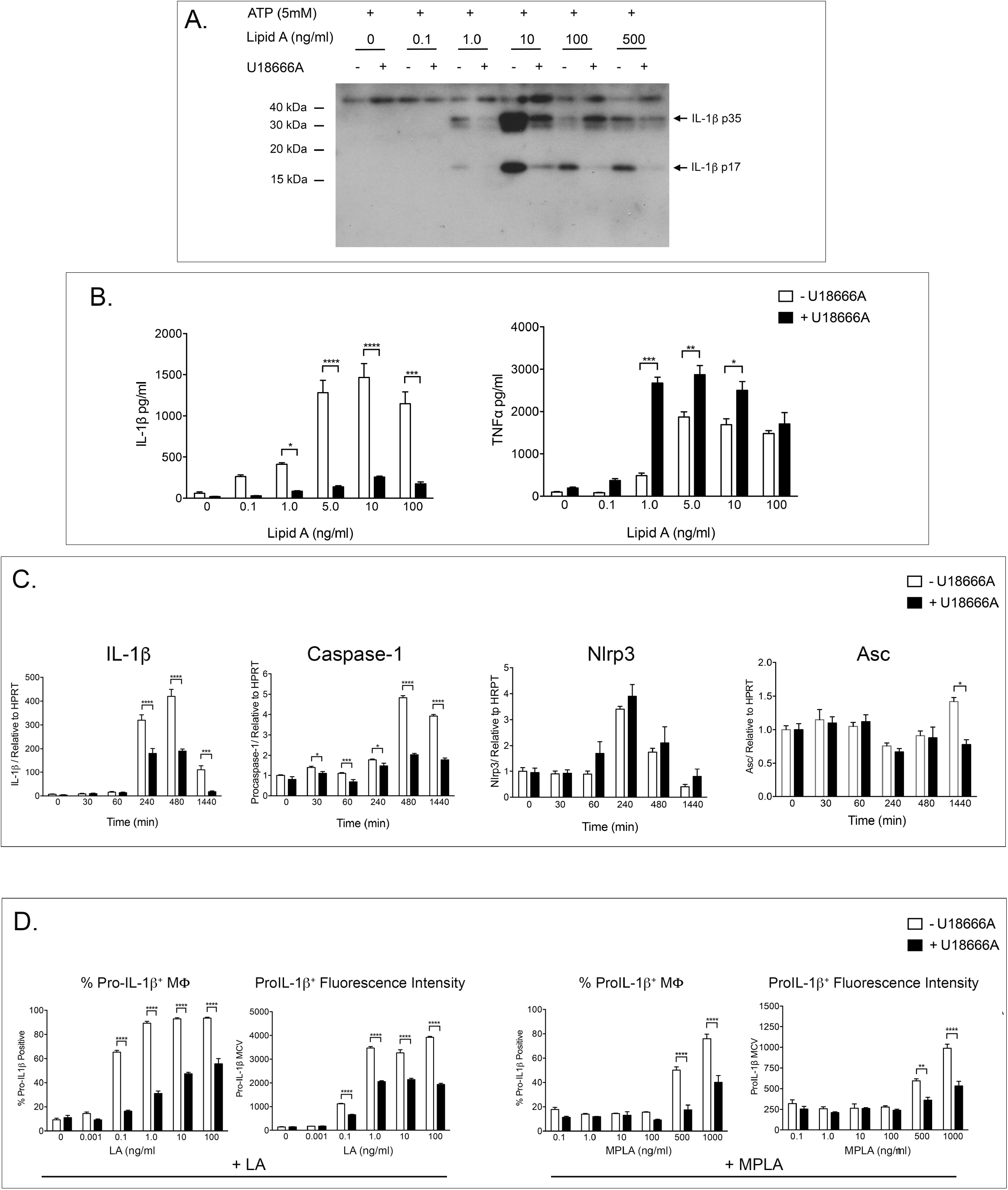
Treatment of RAW 264.7 MΦ with U18666A results in significantly impaired priming of the NLRP3 inflammasome. Panel A. Western blot of culture supernatants of vehicle or U18666A treated RAW 264.7 MΦ primed with various concentrations of LA and activated with 5 mM ATP. Bands corresponding to ProIL-1β p35 and IL-1β p17 proteins are indicated. Panel B. ELISA determinations of IL-1β (left histogram) and TNFα (right histogram) concentrations in culture supernatants of vehicle treated (open columns) or U18666A (filled columns) RAW 264.7 MΦ primed with various concentrations of LA and activated with 5 mM ATP. Data are mean± SEM, n= 5 per treatment. **** *p*< 0.0001, *** *p*<0.001, * *p*<0.05. 2-way ANOVA. Data are representative of two independent experiments. Panel C. Relative expression of IL-1β, capase-1, Nlrp3 and Asc transcripts in vehicle-treated (open columns) or U18666A-treated (filled columns) RAW 264.7 MΦ primed with 100 ng/ml LA at indicated time points. Data are mean± SEM, n= 5 per treatment **** *p*< 0.0001, *** *p*<0.001. Data are representative of two independent experiments. Panel D. FACS determinations of frequencies or fluorescence intensities of ProIL-1β^+^ vehicle-treated (open columns) or U18666A-treated RAW 264.7 MΦ following stimulation with various doses of LA or monophosphoryl LA (MPLA) for 6h. MCV = mean channel value. Data are mean± SEM, n= 5 per treatment. **** *p*< 0.0001, *** *p*<0.001, ** *p*<0.01. 2-way ANOVA. Data are representative of three independent experiments.

To quantify the effect of U18666A-inhibition of Npc1 activity on the generation of ProIL-1β we treated RAW 264.7 MΦ with U18666A and then stimulated with different concentrations of LA or the weaker TLR4 agonist, monophosphorylated LPS^31^. FACS analysis showed a significantly reduced frequency of ProIL-1β^+^ cells and lower staining intensity for cells that had been pre-treated with U18666A and primed with either of the two TLR4 ligands (Fig 4D).

### Reduced ProIL-1β generation by U18666A-treated RAW 264.7 MΦ is not dependent upon lysosomal cholesterol accumulation but is mimicked by drugs that modulate calcium

We exploited the pharmacological model of NPC to investigate the pathological mechanism(s) underlying the lysosomal storage disorder that might be responsible for impaired priming of the NLRP3 inflammasome. NPC is characterized by a complex pathological cascade^13,20,21^. One of the most prominent cellular features of NPC is the accumulation of LDL-derived free cholesterol in the endo-lysosomal system, which was the basis for the clinical diagnosis of this disorder for many years^20^. To enable U18666A-mediated induction of NPC phenotypes without lysosomal cholesterol storage, we grew RAW 264.7 MΦ in lipoprotein-deficient serum (LPDS) for 48h and then treated with U18666A. In comparison to cells grown in medium containing serum (as a source of LDL-cholesterol) and treated with U18666A, filipin staining showed the predicted absence of lysosomal cholesterol accumulation in MΦ from lipoprotein-deficient cultures treated with U18666A (Fig 5A). Cells grown in either fetal bovine serum or LPDS were then treated with vehicle or U18666A, primed with LA and stained and analyzed by FACS for ProIL-1β expression. The same deficit in ProIL-1β generation was observed in cells cultured in either complete or lipoprotein-depleted media with no statistically significant differences in the frequency (Fig 5B left panel) or staining intensity (Fig 5B right panel) of ProIL-1β^+^ RAW 264.7 MΦ, suggesting that storage of cholesterol is not required for the NLRP3 priming defect in NPC cells.

**Fig 5.**
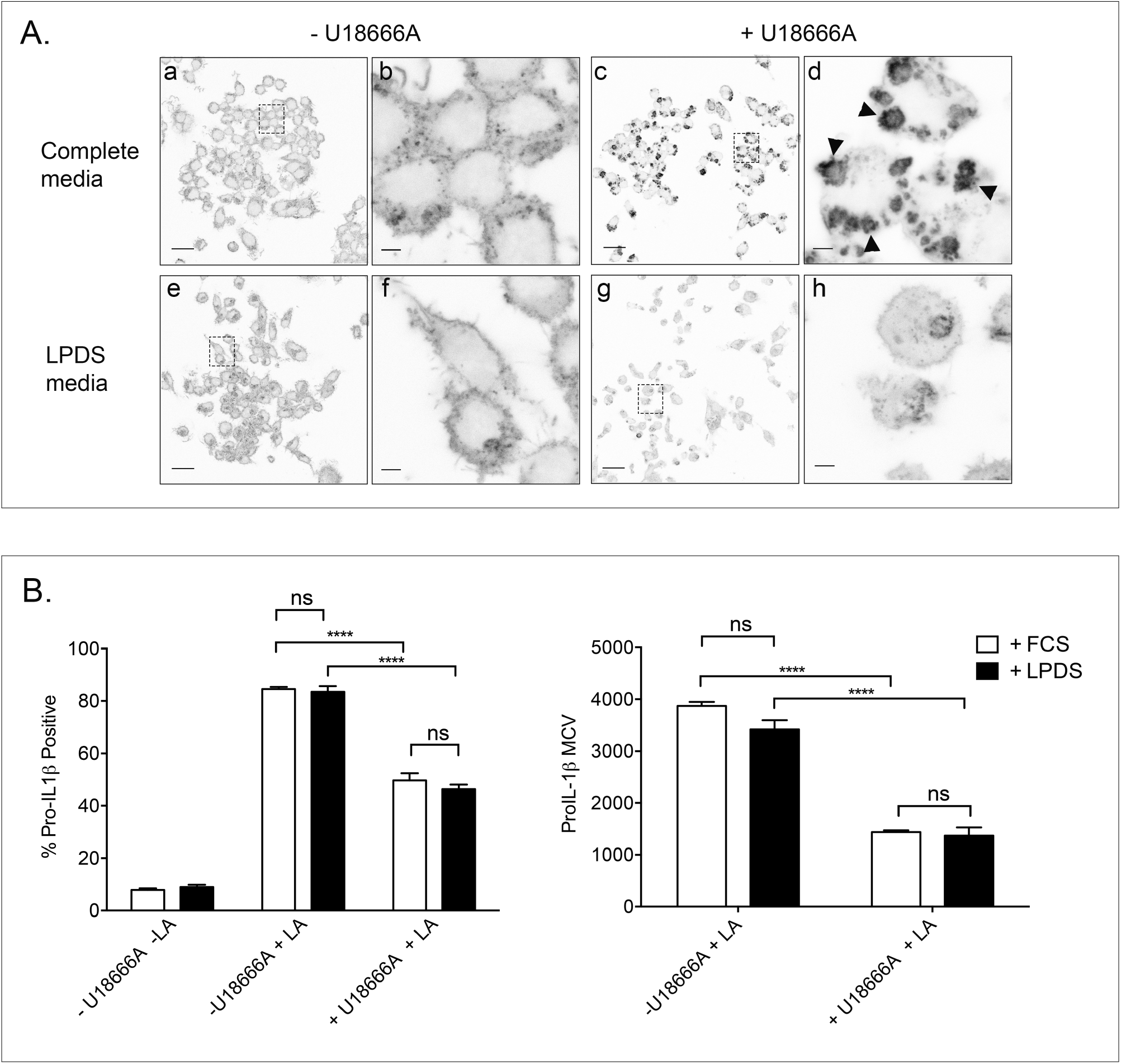
Accumulation of lysosomal cholesterol is not responsible for inhibition of NLRP3 priming by U18666A. Panel A. Representative images of RAW 264.7 MΦ cultured in media containing 10% (v/v) fetal bovine serum – complete medium (a-d) or lipoprotein-deficient serum (LPDS) (e-h) stained with filipin to reveal cholesterol distribution. Images a and b are cells in cultured in complete medium in the presence of vehicle; images c and d are RAW 264.7 MΦ in the same medium and treated with 2 μg/ml U18666A for 24h. Images e and f and g and h are RAW 264.7 MΦ cultured in LPDS, either without or with U18666A. Images b, d, f and h are magnifications of the area identified in the previous image. Scale bars represent 20 μm (a, c, e and g) or 2.5 μm (b, d, f and h). Arrow heads indicate examples of lysosomal cholesterol accumulation. Panel B. Histograms of FACS data showing the frequencies (left histogram) and relative fluorescence intensities (right histogram) of RAW 264.7 MΦ cultured in the presence of either fetal bovine serum (open columns) or LPDS (filled columns), treated with vehicle or 2 μg/ml U18666A and stimulated with 100ng/ml LA. MCV = mean channel value. Data are mean± SEM, n= 5. **** *p*< 0.0001; ns, not significant. 2-way ANOVA. Data are representative of two independent experiments.

We have reported previously that dysfunctional NPC1 results in the accumulation of a broad range of lipids, of which the storage of sphingosine in the lysosome results in a significant reduction in acidic compartment Ca^2+^ levels and loss of divalent cation homeostasis^21^. Failure to release sufficient Ca^2+^ from the lysosome is compatible with the cellular phenotypes of NPC and elevation of cytosolic Ca^2+^ normalizes these phenotypes and provides significant therapeutic benefit in *Npc1*^-/-^ mice^21^. We therefore explored whether pharmacological manipulation of cellular Ca^2+^ stores and in particular lysosomal Ca^2+^ would affect the generation of ProIL-1β.

Pre-treatment of RAW 264.7 with BAPTA-AM, a cell-permeant Ca^2+^ chelator that is highly selective for Ca^2+^ and is largely insensitive to pH^32^ significantly reduced both the frequency (Fig 6A) and staining intensity (Fig 6B) of ProIL-1β^+^ cells. 2-aminoethoxydiphenyl borate (2-APB), a blocker of store-operated Ca^2+^ entry and IP3 receptors^33^ also significantly reduced ProIL-1β production (Fig 6A and B). Finally, exposure to trans Ned-19, a specific antagonist of nicotinic acid adenine dinucleotide phosphate (NAADP) mediated lysosomal Ca^2+^ release^34^ significantly reduced ProIL-1β generation following inflammasome priming (Fig. 6A and B).

**Fig 6.**
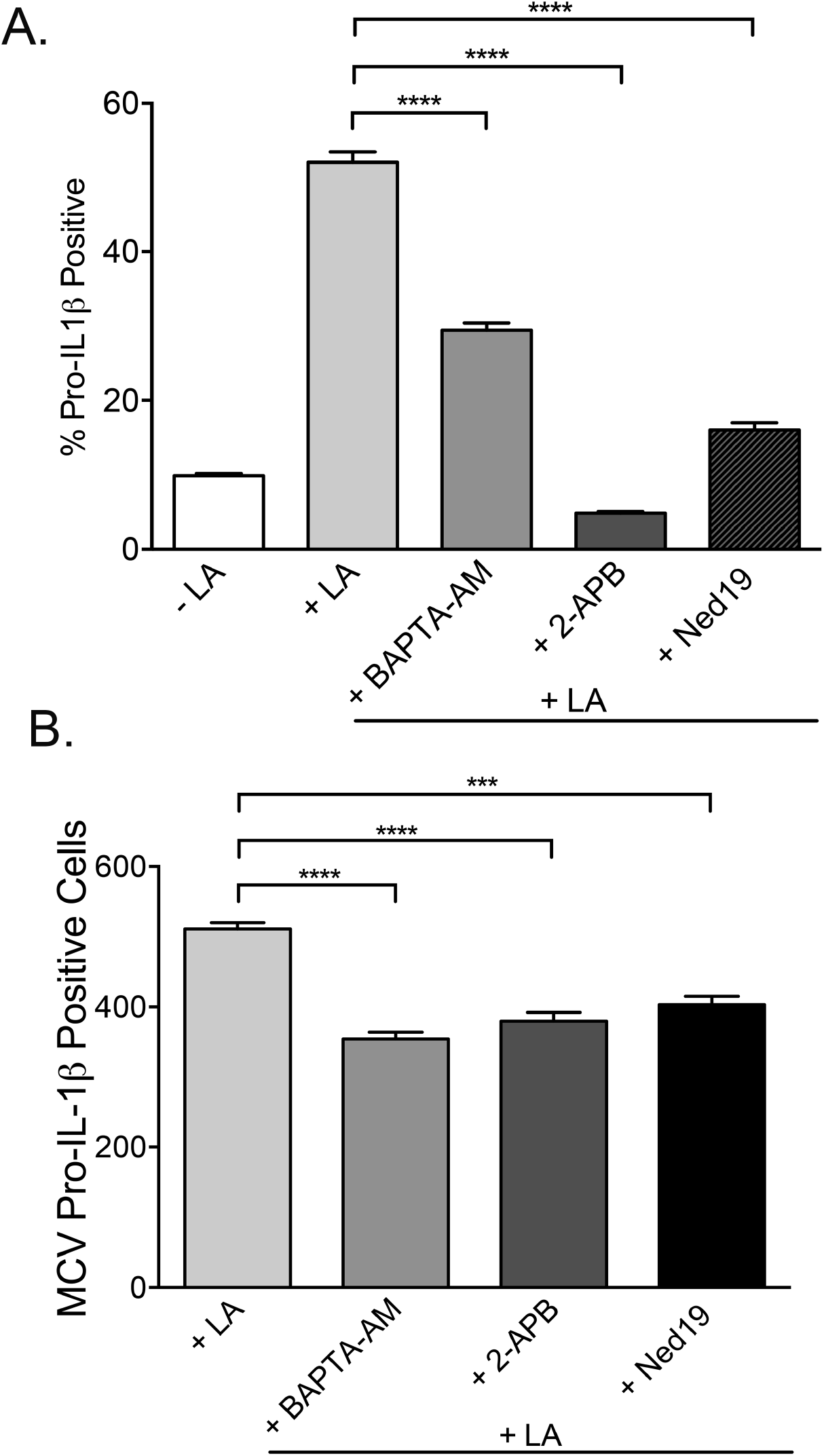
Specific drugs that modulate Ca^2+^ availability from different cellular stores inhibit NLRP3 priming in RAW 264.7 MΦ. Panel A. Frequencies of ProIL-1β^+^ RAW 264.7 MΦ either unstimulated, primed with 100 ng/ml LA for 6h or pre-treated with Ca^2+^ modulating drugs and then primed. Data are mean± SEM, n= 5 per treatment. **** *p*< 0.0001. Student t-test. Data are representative of three independent experiments. Panel B. Relative fluorescence intensities of ProIL-1β^+^ RAW 264.7 MΦ either unstimulated, primed with 100 ng/ml LA for 6h or pre-treated with Ca^2+^ modulating drugs prior to priming. MCV = mean channel value. Data are mean± SEM, n= 5 per treatment. **** *p*< 0.0001, *** *p*<0.001. Student t-test. Data are representative of three independent experiments.

### Loss of the endo-lysosomal Ca^2+^ permeable two-pore channels (Tpcns) Tpcn1 and Tpcn2 significantly impairs ProIL-1β generation

The effects of the Ca^2+^ modulating compounds upon synthesis of ProIL-1β and in particular inhibition by trans Ned-19 suggested a requirement for acid store Ca^2+^ release for efficient priming of the NLRP3 inflammasome. To investigate this possibility directly we took advantage of mice engineered to lack the lysosomal Ca^2+^ release channels, two-pore channel 1 (*Tpcn1^-/-^*) or two-pore channel 2 (*Tpcn2^-/-^*). We prepared MΦ from mice deficient in either Tpcn1 or Tpcn2, the two mammalian members of the Tpcn family of lysosomal channels permeable to Ca^2+^ and investigated their capacity to generate ProIL-1β in response to LA stimulation. Bone marrow-derived macrophages (BM-MΦ) lacking either Tpc1 or Tpc2 showed significantly impaired production of ProIL-1β after priming as compared to wild type cells, with a lower frequency of ProIL-1β^+^ cells (Fig 7A) and reduced staining intensity (Fig 7B). To verify that this deficit was replicated by tissue MΦ, we isolated, LA-stimulated, stained and FACS analyzed *Tpc2*^-/-^ RPMΦ. There was a decreased number of ProIL-1β^+^ *Tpc2*^-/-^ RPMΦ, (Fig 7C), which also displayed lower fluorescence intensity (Fig 7 D).

**Fig 7.**
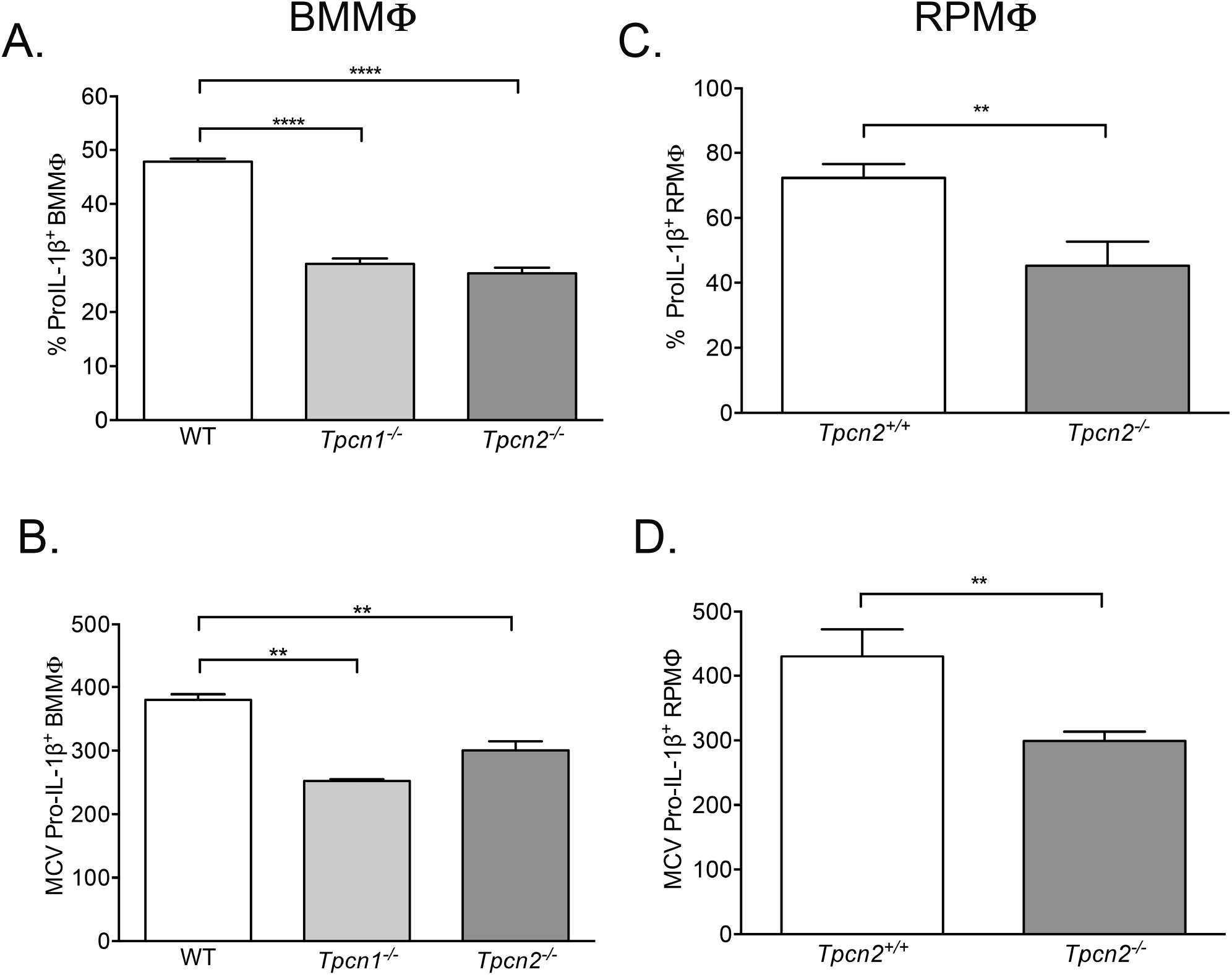
*Tpcn1*^-/-^ and *Tpcn2*^-/-^ BMMΦ and *Tpcn2*^-/-^ RPMΦ display significantly impaired generation of ProIL-1β. Histograms of FACS data of WT, *Tpcn1*^-/-^ and *Tpcn2*^-/-^ BMMΦ (left panels) and WT and *Tpcn2*^-/-^ RPMΦ (right panels) which had been primed with 100ng/ml LA and stained for ProIL-1β. A, frequencies of ProIL-1β positive BMMΦ; B, relative fluorescence intensities of ProIL-1β positive BMMΦ; C, frequencies of ProIL-1β positive RPMΦ and D, relative fluorescence intensities of ProIL-1β positive RPMΦ. MCV = mean channel value. Data are mean± SEM, n= 6. **** *p*< 0.000, ** *p*<0.01. 2-way ANOVA. Data are representative of three independent experiments.

### LA-mediated NLRP3 priming in RPMΦ in Tpc2^-/-^ mice is impaired in vivo

In order to investigate the physiological requirement for lysosomal Ca^2+^ release for efficient NLRP3 priming *in vivo*, we established a low dose LA stimulated sterile peritonitis protocol in *Tpc2*^-/-^ mice that was sufficient to induce expression of ProIL-1β by RPMΦ but did not result in significant loss of the cell population from the peritoneal cavity^35^. We injected 50 ng/kg of LA intraperitoneally (or PBS vehicle), recovered the peritoneal cellular lavage at 3h post-injection and stained with anti-F4/80 and ProIL-1β antibodies and FACS analyzed. There were statistically significantly fewer ProIL-1β^+^ F4/80^hi^ RPMΦ recovered from *Tpc2*^-/-^ animals, which had lower intensity of staining for the cytokine precursor than RPMΦ from *Tpc2*^+/+^ mice (Fig. 8 A and B).

**Fig 8.**
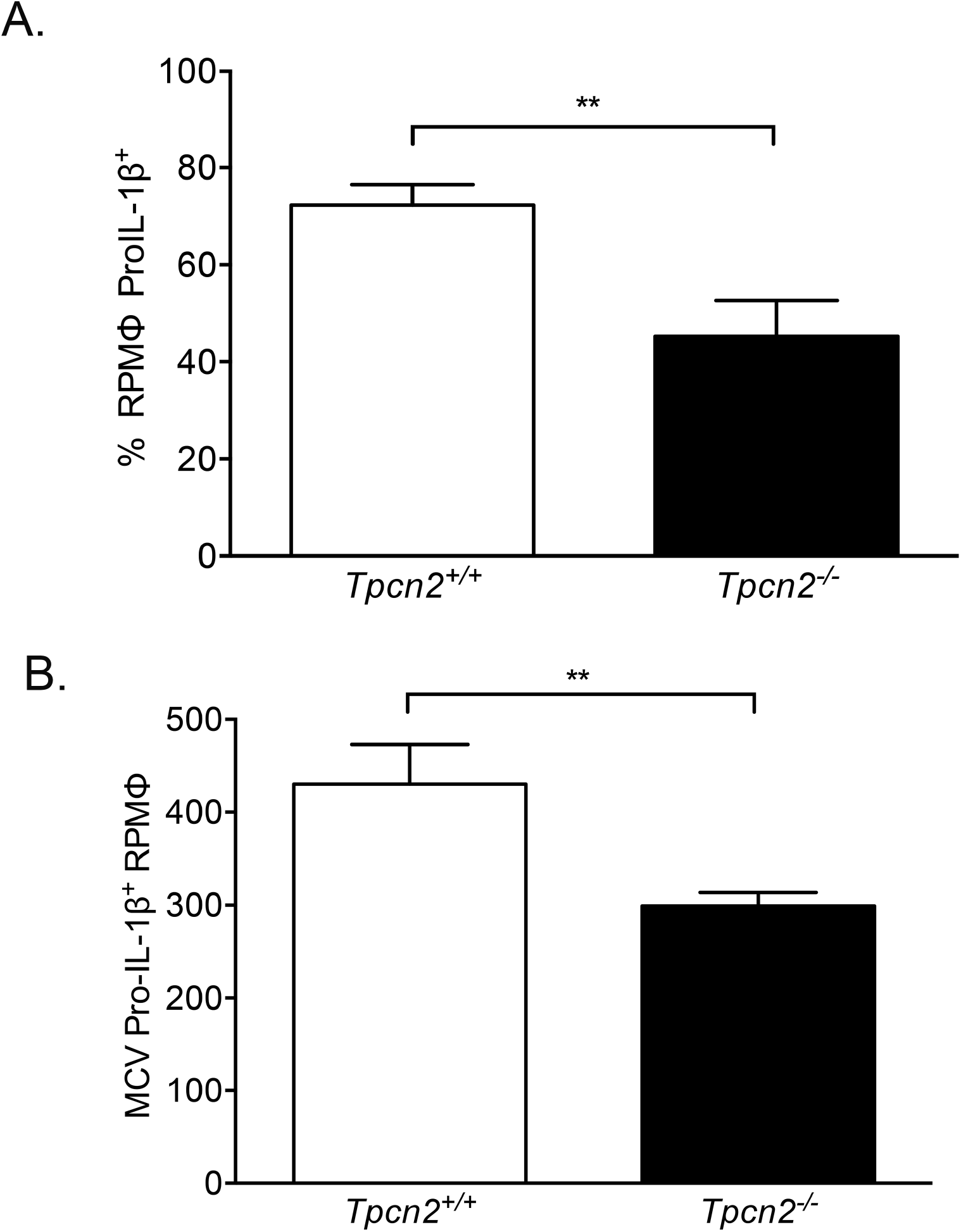
*Tpcn2*^-/-^ RPMΦ display significantly impaired generation of ProIL-1β *in vivo*. Histograms of FACS data of *Tpcn2*^+/+^ and *Tpcn2*^-/-^ RPMΦ recovered from mice 3h after intraperitoneal injection with 50ng/kg LA. A, frequency of ProIL-1β positive RPMΦ; B, relative fluorescence intensity of ProIL-1β positive RPMΦ. MCV = mean channel value. Data are mean± SEM, n= 6. ** *p*<0.01. Student t-test. Data are representative of three independent experiments.

## Discussion

IL-1β is a biologically potent cytokine that induces inflammation and modulates immune responses^23,24,36^. It is therefore not surprising that inflammasome-dependent generation of IL-1β has evolved to be tightly regulated at multiple levels^25,37^. Canonical production of IL-1β by the most studies inflammasome, NLRP3, requires two signals^37,38^. An initial priming event resulting from ligand binding of pattern recognition or other receptors engages the NF-kB signal transduction pathway, resulting in the transcription and translation of ProIL-1β and other inflammasome components such as NLRP3^39^. The second step is inflammasome activation that involves inflammasome assembly, caspase-dependent processing of the cytokine precursor pro-IL-1β and the subsequent secretion of mature IL-1β ^37^. NLRP3 activation can be caused by diverse stimuli, implying the existence of a shared downstream mechanism. Understanding the orchestration of this complex cellular machinery that constitute this activation step has been a primary research focus^27–40^. In contrast, the regulation of the priming step has been much less well studied and therefore remains incompletely understood. Here, from studying a rare inborn error of metabolism, the lysosomal storage disease NPC, we have discovered that the priming of the NLRP3 inflammasome is also subject to regulation and we show for the first time that ProIL-1β synthesis requires Ca^2+^ release specifically from acidic lysosomal stores.

Although the occurrence of aberrant pro-inflammatory responses is widespread amongst LSDs, indicating that loss of lysosomal homeostasis can readily provoke inflammation, current understanding of their pathophysiology is incomplete. Because studies made in other LSDs such as Gaucher disease have reported increased IL-1β secretion^41^ we were surprised to observe significantly reduced production of this cytokine specifically in NPC, whereas generation of other pro-inflammatory cytokines such as TNFα was unchanged in NPC macrophages. We interpret these data to indicate that disease-specific alterations in lysosomal properties determine the nature of the altered inflammatory profile in LSDs and that the lysosome is involved in the generation of specific inflammatory responses. Furthermore, analysis of the impact of loss of Npc1 activity upon NLRP3-dependent IL-1β production using different methodologies, both genetic and pharmacological, verified that the deficit was due to impairment of the initial priming step and that NLRP3 inflammasome activation was not apparently affected.

In order to determine whether reduced generation of IL-1β might be indicative of a more generalized inhibition of the pro-inflammatory response by NPC MΦ, we analyzed secretion of TNFα which is also stimulated by TLR4 ligands^42^. Interestingly, for RPMΦ there was no significant difference between production of TNFα by *Npc1*^+/+^ and *Npc1*^-/-^ cells, but in the case of U18666A-treated RAW 264.7 MΦ, which also showed impaired IL-1β production, we measured significantly greater TNFα secretion by drug treated cells. We cannot fully explain this difference but it is possible that the relative MΦ activation state may influence the magnitude of other components of the pro-inflammatory response when Npc1 is inactivated either genetically or pharmacologically.

Lysosomes are recognized to be involved in NLRP3 inflammasome-dependent IL-1β production, but the evidence is for affecting activation, not the priming step^43^. Several studies have shown that an insult to the lysosome, such as physical disruption caused by phagocytosis of particulates results in activation of the inflammasome in primed cells. For example, phagocytosis of silica particles causes lysosome destabilization and membrane permeabilization which results in K^+^ efflux and inflammasome activation^44,45^, as do alum-containing adjuvants^46^. Loss of lysosomal acidification through inhibition of the vacuolar H^+^-ATPase with bafilomycin prevents K^+^ efflux across the plasma membrane and NLRP3 activation by particulate matter ^47^. The relative extent of lysosomal membrane disruption may affect the degree of NLRP3 activation ^48^. However, as far as we are aware the lysosome has not been linked previously to priming of the NLRP3 inflammasome.

We reasoned that the impaired NLRP3 inflammasome priming we observed in NPC cells would result from one or more of the cellular phenotypes of this specific lysosomal disorder. NPC is defined by a complex pattern of cellular phenotypes that include accumulation of multiple lipid species (cholesterol and sphingolipids), inhibition of trafficking events in the endo-lysosome system and loss of lysosome: ER contact sites^21,20,30^. Indeed, this complexity has made demarcation of the pathological cascade underlying NPC challenging and subject to differential interpretation^49^. Lysosomal storage of LDL derived free cholesterol is a prominent feature in NPC. However, we found that culturing RAW 264.7 MΦ in media containing lipoprotein-deficient serum significantly reduced U18666A-dependent lysosomal cholesterol accumulation but did not impact significantly upon inhibition of LA-stimulated ProIL-1β expression, indicating that impairment of NLRP3 priming was unlikely to be the result of storage of cholesterol. It is interesting to note that our findings of reduced NLRP3-dependent generation of IL-1β has been reported in a different model of NPC ^50^. In contrast to the studies shown here in which primary MΦ obtained from an authentic mouse model of the storage disease were analyzed *in vitro* and *in vivo*, the published investigation focused primarily on the use of an immortalized BMMΦ cell line in which mutations in Npc1 had been engineered by CRISPR-cas9 technology. The authors provided evidence of abrogated NLRP3 inflammasome activation in the system with no apparent changes to NLRP3 inflammasome priming and concluded that loss of cholesterol organelle homeostasis and perturbed trafficking was responsible for the phenotype. Although at this time we cannot fully reconcile all of the differences between the two studies, both studies reported changes to NLRP3 activity that result in significantly lower secretion of IL-1β. A relationship between cholesterol biosynthesis and metabolic signaling and NLRP3 inflammasome activation has been broadly recognized; however, the precise mechanistic basis is less clear because of evidence of the direct effect of the master regulator SREBP2 on the escort activity of SCAP affecting NLRP3 inflammasome activation rather than indirectly via changes to cholesterol homeostasis^51^. The fact that we see the priming defect in pharmacological and genetic models of NPC and in Tpcn deficient mice further support a defect in priming as a major factor in reduced IL-1β secretion in Npc1 deficient cells.

Our analysis of NLRP3 priming in NPC cells included determination of relative abundance of relevant transcripts by Q-PCR. We confirmed significantly reduced IL-1β and caspase-1 transcription in LA primed NPC MΦ, but unexpectedly, the profile of Nlrp3 transcription after stimulation with LA was not different from controls. Bauernfeind and colleagues ^39^ have reported that NF-kB-dependent signals regulate transcription of Nlrp3 and ProIL-1β but not Asc mRNAs in primed MΦ and that expression of NLRP3 is required for NLRP3 activation. The data we present here suggest existence of additional regulatory mechanism(s) that differentially impact gene expression during priming of the NLRP3 inflammasome.

Previously, we reported that NPC1-mutant cells and RAW 264.7 MΦ in which the NPC disease cellular phenotype was induced with U1866A have a significant reduction in the acidic compartment Ca^2+^ store (to approximately 25-30% of wild type levels) as a consequence of sphingosine accumulation^21^. Evidence supports the hypothesis that sphingosine building up in the lysosome is an initiating factor in NPC disease ^22,52^ and it was confirmed that compensation for the lack of sufficient Ca^2+^ release by elevating cytosolic Ca^2+^ corrected cellular disease phenotypes and prolonged survival of *Npc1*^-/-^ mice^21^. We therefore investigated whether pharmacological manipulation of Ca^2+^ would affect NLRP3 priming and observed that each of the compounds tested, which were selected to delineate between different sources of Ca^2+^, reduced production of ProIL-1β. Importantly, trans Ned-19, an antagonist of NAADP-stimulated lysosomal Ca^2+^ release via Tpcns^34^ significantly inhibited NLRP3 priming. We further confirmed the involvement of Tpcn-mediated acidic store Ca^2+^ release by determining the extent of NLRP3 priming in MΦ genetically lacking either Tpcn1 or Tpcn2 and found significant inhibition of ProIL-1β production when either channel was deficient. Expression of Tpcn2 is considered to be primarily lysosomal, whereas that of Tpcn1 is broadly expressed in endosomal pathways ^53,54^, suggesting that both cellular compartments may contribute to NLRP3 inflammasome priming. Finally, we demonstrated that LA-mediated priming of *Tpcn2*^-/-^ RPMΦ *in vivo* also resulted in significantly lower synthesis of ProIL-1β, confirming its physiological relevance.

Several investigations of the requirement for Ca^2+^ mobilization in the generation of IL-1β have been reported and a consensus of the effects of abolishing Ca^2+^ mobilization has emerged, but these studies have focused primarily upon the role of the cation in inflammasome activation not the priming step^55^. Brough *et al* ^56^ showed that the permeant chelator BAPTA-AM prevented IL-1β secretion by LPS primed, ATP activated murine RPMΦ. Ca^2+^ flux and NLRP3 activation mediated by ATP, nigericin and alum were attenuated in MΦ in Ca^2+^ free media or by exposure to thapsigargin, an inhibitor of sarcoplasmic/ER Ca^2+-^ATPase and by 2-APB. These findings are consistent with the requirement for Ca^2+^ release from the endoplasmic reticulum and store-operated extracellular entry of Ca^2+ 57^. However, the effect of extracellular Ca^2+^ has been shown to be complex and the effects dependent upon concentration. In the absence of other activation signals Ca^2+^ alone triggered IL1-β secretion by LPS-primed BMDMs but it inhibited ATP-driven IL-1β release through mechanisms that required the Ca^2+^-sensing receptor, CASR ^58^. LPS-induced gene expression is also dependent upon the chanzyme, TRPM7, a non-selective Ca^2+^-conducting ion channel, which is responsible for cation influx necessary for TLR4 endocytosis^59^. Although IL-1β transcription in LPS stimulated *Trpm7*^-/-^ BMMΦ was significantly reduced in comparison to wild type cells, similar to what we report here, but unlike *Npc1*^-/-^ RPMΦ there was also a significant reduction in other pro-inflammatory cytokines including TNFα and IL-6 ^59^.

The Ca^2+^ content of late endosomes/lysosomes is not large due to the relatively small size of the acidic compartment and this organelle does not signal globally in the way that, for example the ER does. Ca^2+^ release from the lysosome may be triggered specifically by the second messengers NAADP ^60^ thereby providing one level of regulation which results in the generation of localized Ca^2+^ signals. Bi-directional communication between the lysosome and ER means these organellar interactions can be amplified or tempered to modulate cytosolic Ca^2+^ levels^11,61,62^ and thus potentially have effects more globally by lysosomal Ca^2+^ release triggering Ca^2+^ induced Ca^2+^ release from the ER^61^. There is evidence for the importance of signaling through lysosomes and late endosomes in an increasing number of distinct cellular processes^63^. For example, lysosomal Ca^2+^ release has been shown to modulate the function of the immune system as illustrated by the requirement for exocytosis of T cell cytolytic granules^64^, expulsion of exosome-encased bacteria^65^, remodeling of NK cell secretory lysosomes^66^ and viral infection^67^. It is especially relevant that mobilization of lysosomal Ca^2+^ has been demonstrated in the innate immune processes of phagocytosis^68^, phagosome maturation^69^ and ion fluxes in the endo-lysosome necessary for tissue surveillance by MΦ ^70^. In light of the data provided here on priming of the inflammasome, we speculate that release of acidic store Ca^2+^ may serve to coordinate triggering of multiple immune cell activities after engagement with specific microbes.

In summary, acidic store Ca^2+^ mobilization is required for efficient priming of the NLRP3 inflammasome and thereby provides an additional level for the control of cytokine production and has the potential to be a novel target for anti-inflammatory intervention.

## Materials and Methods

### Animals

Breeding colonies of BALBc/NPC^NIH^ (*Npc1^m1n^*), here referred to as *Npc1*^-/-^ ^71^, *Tpcn1^tm1Dgen^* (*Tpcn1^-/-^*) and *Tpcn2^Gt9YHD437)Byg^* (*Tpcn2^-/-^*) ^72^ mice were maintained at the University of Oxford under specific pathogen free housing. All animal use was approved under the authority of a license issued by the UK Home Office (Animals [Scientific procedures] Act 1986).

### Cells and culture

RAW 264.7 and J774. A murine MΦ were obtained from ATCC and maintained in RPMI 1640 supplemented with 10% (v/v) fetal calf serum, 1% glutamate, 1% penicillinstreptomycin (complete media) and passaged at regular intervals to ensure viability was > 90%. Resident peritoneal MΦ (RPMΦ) were harvested from age matched wild type and symptomatic *Npc1*^-/-^ mice and from 8–12-week-old wild type and *Tpcn2*^-/-^ mice by peritoneal lavage, plated out in complete media and allowed to adhere and non-adherent cells washed off. RPMΦ were allow to rest overnight before use. For the generation of bone marrow-derived MΦ (BMMΦ) bone marrow was flushed from the tibia and fibula bones and resuspended in complete medium containing 30% (v/v) L929 cell-conditioned media as a source of M-CSF ^73^. Media was replaced with fresh media on days 3 and 6.

Cells were stimulated with either synthetic lipid A (Kdo2-Lipid A, AdipoGen), peptidoglycan (Invivogen) or monophosphoryl lipid A (Invivogen) at the indicated concentrations. When required, primed cells were washed and then activated with 5 mM ATP diluted in Opti-MEM media (Thermo Fisher) for 60 min and supernatants collected.

NPC cellular phenotypes were induced in RAW 264.7 and J774.A MΦ by treating with 2 μg/ml U18666A (Merck) for 30h or 72h respectively. To pharmacologically modulate Ca^2+^ availability from cellular stores, cells were pre-treated with either BAPTA-AM, 2-aminoethoxydiphenyl borate (2-APB) or trans-Ned-19 (all Tocris).

### Flow cytometry (FACS)

Live cell samples were blocked with FC block (BD Biosciences) prior to staining with titrated anti-F4/80-APC antibody (Biolegend) on ice and then washed. Where indicated, cells were then stained with LysoTracker™green DND-26 according to te Vruchte *et al* ^28^ and immediately analyzed. For determination of ProIL-1b expression, cells were fixed and permeabilized using fixation and permeabilization/wash buffers (Biolegend) according to the manufacturer’s instructions and then stained with anti-murine ProIL-1β antibody (clone NJTEN3, Thermofisher), washed and analyzed. All samples were acquired on a FACSCanto II (BD Biosciences) which had been set up using BD 7 colour setup beads. A minimum of 1 × 10^4^ events were collected for each sample. Data were gated and analyzed using FlowJo software (FlowJo, LLC).

### ELISAs

Cytokine concentrations in culture supernatants were determined using specific ELISA kits (IL-1β and TNFα, Thermo Fisher Scientific) according to instructions provided by the manufacturer.

### Q-PCR

RNA from MΦ was isolated using Monarch total RNA MiniPrep kit (BioLabs) according to the manufacturer’s protocol. cDNA was synthesized using iScript cDNA synthesis kit (BIORAD). Reaction mixes containing cDNA template, PowerUp SYBER Green (ThermoFisher) and specific primers were run on a CFX96 Real-Time PCR system (BIO-RAD). Assays specific for relative expression of murine β-actin was used to normalize for input.

### Western blotting

Cell lysates were prepared using cell lysis buffer (Cell Signaling), incubated on ice for 60 min and spun at 4°C at 13K to remove insoluble material. Protein concentrations of lysates were determined using BCA assay (Merck). Proteins in culture supernatants were precipitated using methanol/chloroform, the pellet briefly dried and re-suspended in sample loading buffer. Protein samples were diluted with SDS-PAGE reducing gel loading buffer (Thermofisher) and heated at 95°C for 5 min. Samples were loaded onto either 15% or 4-12% SDS-PAGE gels (Thermofisher). Gels were transferred to PVDF membrane using a Trans-Blot Turbo Transfer system (Bio-rad) and membranes blocked with 5% non-dairy milk powder in tris-buffered saline containing 0.1% Tween 20, incubated with anti-IL-1b antibody (R&D Systems) overnight at 4°C, washed and incubated with anti-goat HRP secondary antibody (Jackson Immunoresearch), washed and developed with SuperSignal™ West Pico Plus Chemiluminescent substrate (Thermo Scientific). Gel images were collected on a ChemiDoc XRS+ Imaging System (Bio-rad) and images processed using Imagelab software (Bio-rad). To validate equivalent protein loading in each gel lane where appropriate, membranes were re-probed with anti-β-actin HRP-conjugated antibody (Merck), washed and developed using Pierce™ ECL Western Blotting Chemiluminescent substrate (Thermo Scientific). Images were collected and processed as described above.

## Acknowledgements and funding

We would like to thank Dave Smith and Claire Smith for animal husbandry and genotyping, staff of BSU, University of Oxford for animal maintenance and to all members of the Platt laboratory for helpful discussions. This work was funded by an Investigator in Science Award from The Wellcome Trust to FMP.

---

**Supplementary Fig 1.**
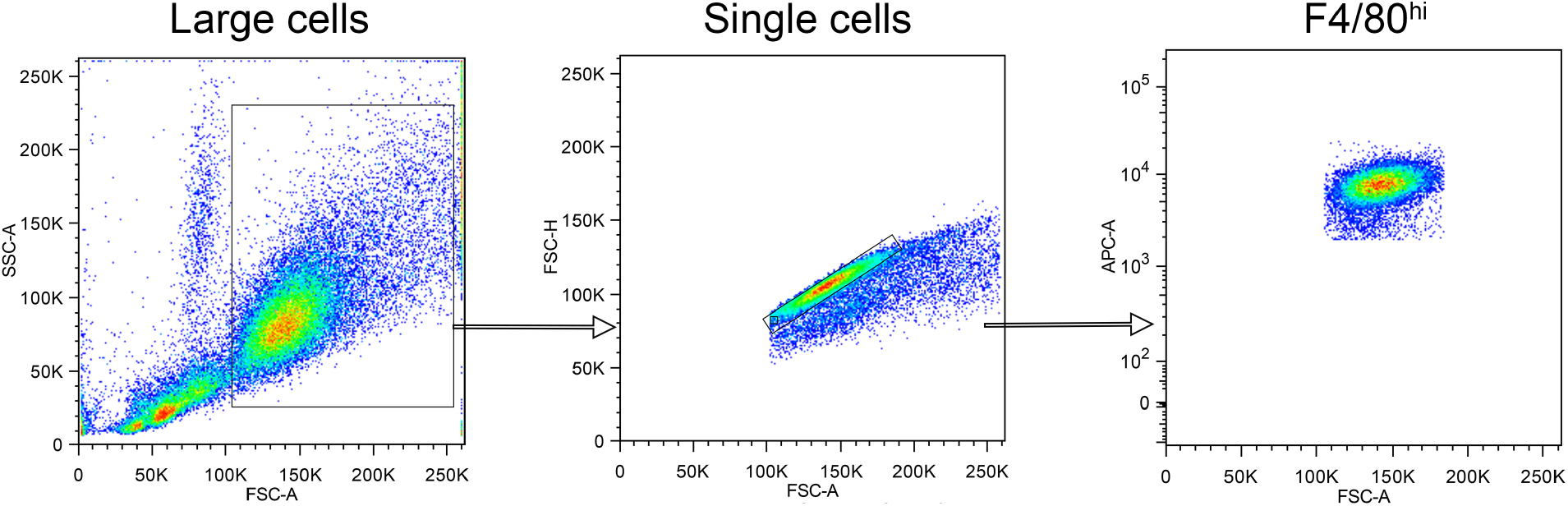
FACS gating strategy for identification of RPMΦ. Cells were initially gated based upon their relative size (FSC-A), single cells were then identified and subsequently F4/80^hi^ RPMF identified.

**Supplementary Fig 2.**
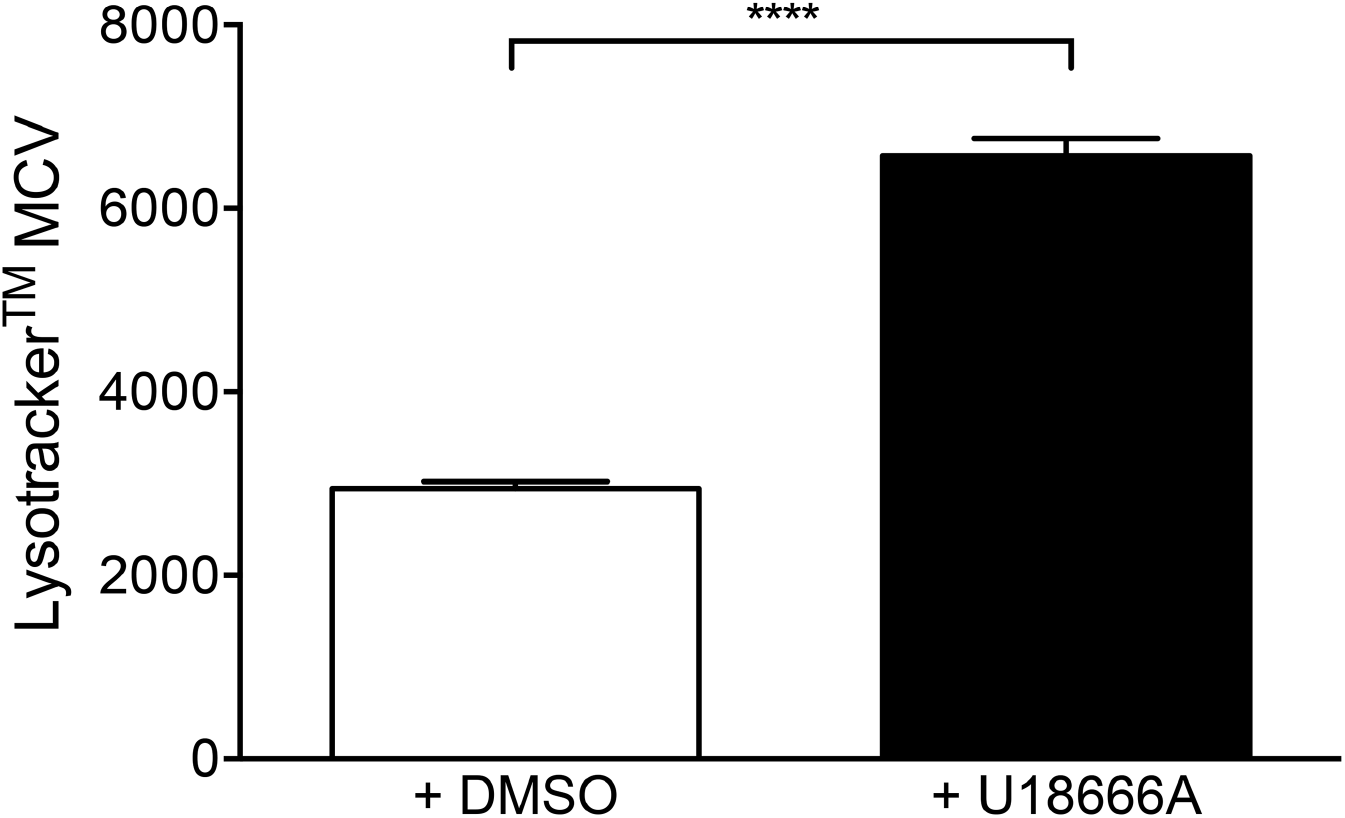
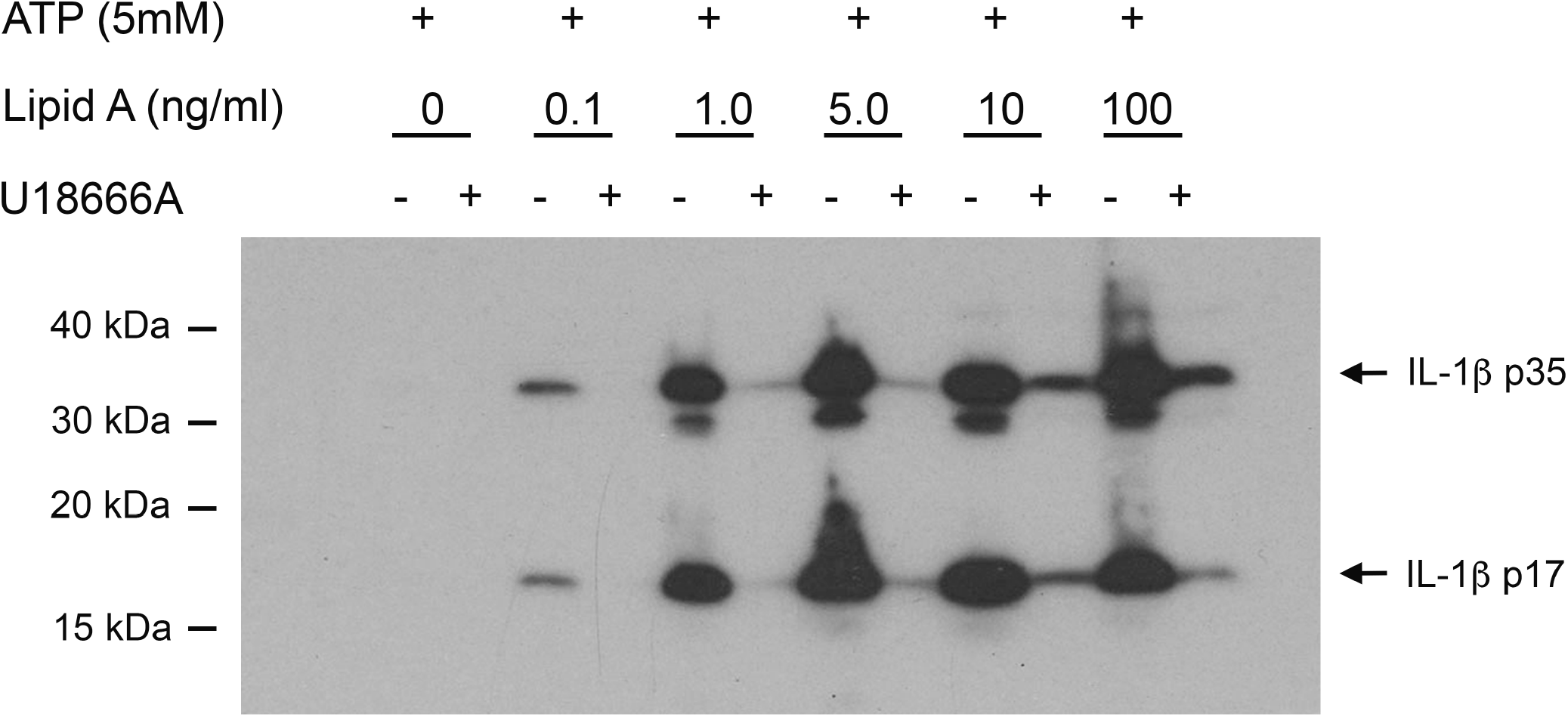
Treatment with U18666A significantly increases LysoTracker™ staining of RAW 264.7 MΦ. Histogram of FACS data of relative fluorescent intensity (MCV) of RAW 264.7 MΦ treated with vehicle or 2 μg/ml U18666A for 30h and stained with LysoTracker™ green DND-26. Data shown are mean± SEM, n= 6. **** *p*< 0.0001. Data are representative of a minimum of six independent experiments.

**Supplementary Fig 3. Treatment with U18666A significantly decreases IL-1 secretion by primed and activated J774A MΦ.** Western blot of culture supernatants of J774 cells primed with different concentrations of LA and then activated with 5 mM ATP separated on SDS-PAGE gel, transferred and probed for IL-1β expression. Arrows indicate bands corresponding to ProIL-1β p35 and IL-1β p17. Gel is representative of two independent experiments.

